# FOXO1 and FOXO3 cooperatively regulate innate lymphoid cell development

**DOI:** 10.1101/2021.01.13.426512

**Authors:** Thuy T. Luu, Jonas Nørskov Søndergaard, Lucía Peña-Pérez, Shabnam Kharazi, Aleksandra Krstic, Stephan Meinke, Laurent Schmied, Nicolai Frengen, Yaser Heshmati, Marcin Kierczak, Thibault Bouderlique, Arnika Kathleen Wagner, Charlotte Gustafsson, Benedict J. Chambers, Adnane Achour, Claudia Kutter, Petter Höglund, Robert Månsson, Nadir Kadri

## Abstract

The natural killer (NK) and non-cytotoxic innate lymphoid cells (ILC) lineages play vital role in the regulation of the immune system. Yet understanding of mechanisms controlling NK/ILC development remains incomplete. The evolutionary conserved FOXO family of forkhead transcription factors are critical regulators of cellular processes. We found that the loss of FOXO1 and FOXO3 together caused impaired activation of the NK gene expression program and reduced ETS binding already at the common lymphoid progenitor (CLP) level and a block at the ILC progenitor (ILCP) to NK progenitor transition. FOXO controlled NK cell maturation in organ specific manner and their ability to respond to IL-15. At the ILCP level, disruption of the ILC lineage specific gene programs was associated with broad perturbation of the generation of the non-cytotoxic ILC subsets. We concluded that FOXO1 and FOXO3 cooperatively regulate ILC lineage specification at the progenitor level as well as the generation of mature ILCs.

## INTRODUCTION

The evolutionarily conserved forkhead box transcription factors of the O class (FOXO) are critical regulators of metabolism, lifespan, fertility, proliferation and cellular differentiation (Lin, Dorman et al. 1997, Ouyang and Li 2011). In mammals, the FOXO family is comprised of four members (FOXO1, FOXO3, FOXO4 and FOXO6) that, with the exception of FOXO6, are widely co-expressed throughout the immune system. Growth factor and survival signals activate the phosphoinositide 3-kinase-Akt signaling pathway, which leads to phosphorylation of the FOXOs and their subsequent nuclear exclusion and degradation (Salih and Brunet 2008). This counteracting the FOXO families role in promoting apoptosis (Brunet, Bonni et al. 1999, Dijkers, Medema et al. 2000) and cell cycle arrest (Alvarez, Martinez et al. 2001, Martinez-Gac, Marques et al. 2004). In the adaptive immune system, FOXOs control a wide range of functions including homing and survival of naïve T cells (Kerdiles, Beisner et al. 2009, Ouyang, Beckett et al. 2009, Ouyang and Li 2011), expansion of CD8^+^ memory T cells (Rao, Li et al. 2012, Tzelepis, Joseph et al. 2013), differentiation of regulatory T cells (Harada, Harada et al. 2010, Kerdiles, Stone et al. 2010, Ouyang, Beckett et al. 2010, Ouyang, Liao et al. 2012) as well as B cell lineage commitment, homing and germinal center proliferation (Dengler, Baracho et al. 2008, Mansson, Welinder et al. 2012, Inoue, Shinnakasu et al. 2017).

NK cells are innate immune cells important for controlling viral infection and cancer (Hoglund and Brodin 2010, Kadri, Thanh et al. 2015, Kadri, Wagner et al. 2016). The IL-15 dependent NK cell lineage (Wu, Tian et al. 2017) is – similar to B and T cells - derived from common lymphoid progenitors (CLP) (Fathman, Bhattacharya et al. 2011). Downstream of the CLP, NK cell develop via two hierarchically related NK progenitor (NKP) stages originally defined by the loss of FMS tyrosine kinase 3 (FLT3) on pre-NKPs and the subsequent acquisition of IL-15Rβ (CD122) on refined NKPs (Fathman, Bhattacharya et al. 2011). Recent studies have refined this developmental scheme and through the use of polychromatic reporter mice demonstrated that the pre-NKP compartment represent a heterogeneous population of innate lymphoid cell progenitors (ILCPs) that give rise not only to NK cells but also the non-cytotoxic ILC subsets (Constantinides, McDonald et al. 2014, Walker, Clark et al. 2019, Xu, Cherrier et al. 2019). For simplicity, we hereafter refer to the heterogenous pre-NKP compartment as ILCPs and the refined NKPs as NKPs.

Despite the identification of these intermediate NK cell progenitors and committed ILC progenitors within the ILCP (Walker + Xu), the precise stages in which ILC lineage-specific developmental programs are enabled as well as and the underlying mechanisms that leads to NK lineage restriction remain to be understood in detail. However, on the gene regulatory level several transcription factors including ETS1, NFIL3 and TCF7 have been shown to impact the development of NK cells at the ILCP and NKP progenitor level (Ramirez, Chandler et al. 2012, Male, Nisoli et al. 2014, Goh and Huntington 2017, Jeevan-Raj, Gehrig et al. 2017).

Little is known about the role of the FOXOs in the development of non-cytotoxic ILCs. However, their role in NK cell maturation has been addressed providing contradictory results. Relying on the specific Cre mediated deletion in Ncr1^+^ (NKp46) cells, Deng *et al.* observed a more mature phenotype in NK cells lacking FOXO1 or FOXO1 and 3 while this was not observed in NK cells lacking FOXO3 alone which lead to the conclusion that FOXO1 is dispensable for NK cell development but negatively regulates NK cell maturation (Deng, Kerdiles et al. 2015). Using a similar model, Wang *et al*., in contrast observed that NK cell development was abrogated by the loss of FOXO1 (Wang, Xia et al. 2016). Noteworthy, Ncr1 expression is acquired only after commitment to NK cell lineage and therefore, the reported results do not address a potential role of the FOXOs in early NK cell development. In line with this, deletion of FOXO1 in the hematopoietic stem cell has been reported to resulted in increased frequencies of NK cell progenitors and committed NK cells (Huang, Wang et al. 2019). Together, this underlines the need for further studying the role of the FOXOs in NK cell progenitors and NK cell maturation.

Using ablation of FOXO1 and/or FOXO3 throughout the hematopoietic system, we here show that the FOXOs are critical for NK progenitor development, establishment of the early NK gene regulatory network and NK maturation. In addition, we show that the loss of FOXO perturbs the development of the non-cytotoxic ILC lineages. These findings provide novel insights into NK and ILC development and the gene regulatory program that underpin ILC development.

## RESULTS

### The NK gene expression program is initiated in ILCPs and NKPs

To characterize gene regulation in early ILC and NK cell development, we performed RNA sequencing on FACS sorted Ly6D^neg^ CLPs (Inlay, Bhattacharya et al. 2009, Mansson, Zandi et al. 2010), ILCPs and NKPs (Fathman, Bhattacharya et al. 2011, Constantinides, McDonald et al. 2014, Walker, Clark et al. 2019, Xu, Cherrier et al. 2019) **(Fig. 1A and Supplementary Table 1)**. In agreement with ILCPs representing a developmental stage between CLPs and NKPs (Fathman, Bhattacharya et al. 2011), principal component analysis (PCA) revealed three distinct groups with the first component (PC1 60%) positioning the related progenitor subsets in the expected hierarchical order **(Fig. 1B)**. Further in line with this, ILCPs expressed genes otherwise only expressed (≥0.3 TPM in all replicas) in CLPs or NKPs **(Fig. 1C)**.

**Fig. 1.**
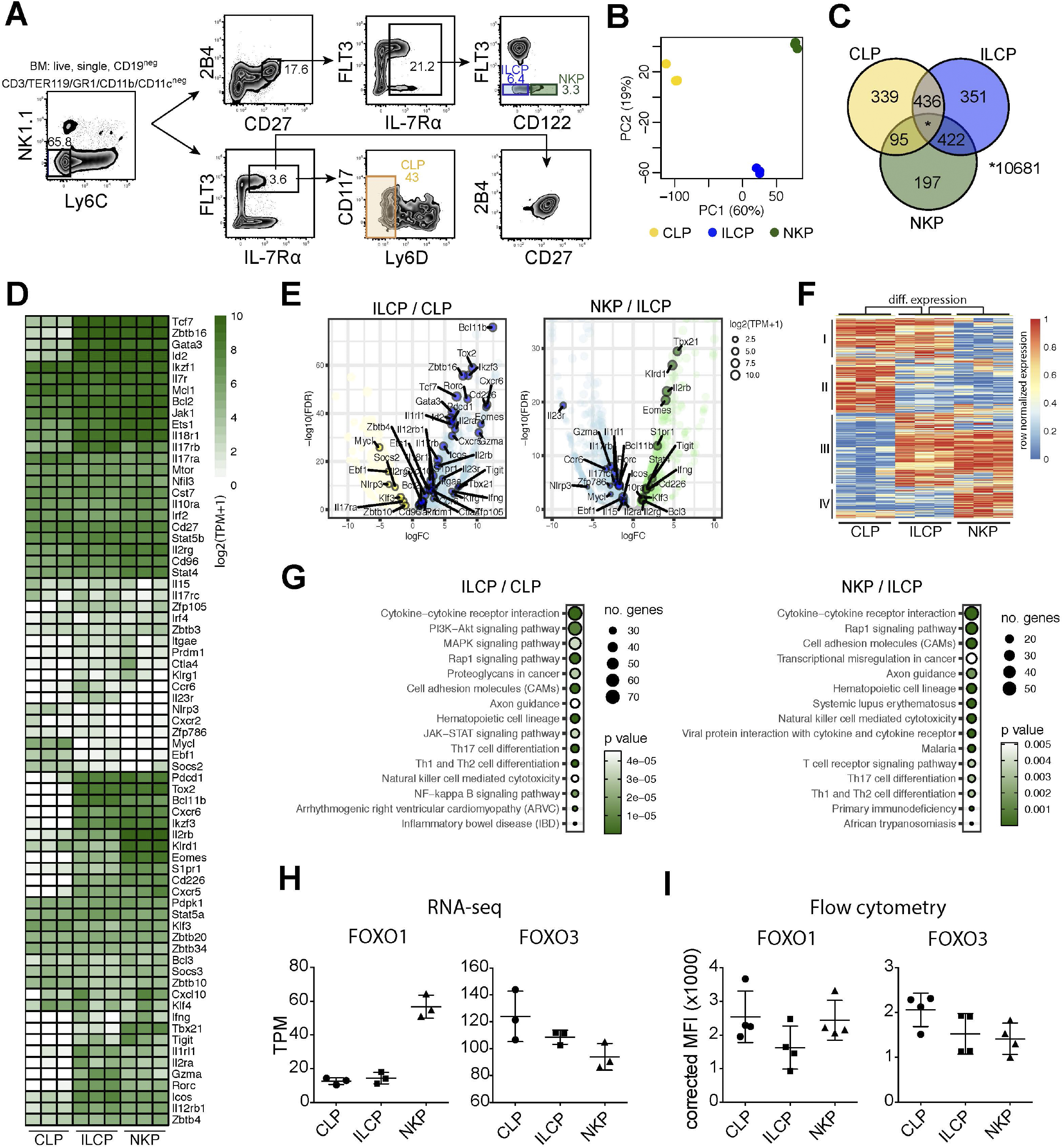
The NKP gene expression program is established gradually in the CLP to ILCP and ILCP to NKP developmental transitions. (A) Gating strategy for FACS sorting of BM NK cell progenitors (Ly6D^neg^ CLP, ILCP and NKP). (B) Principal component (PC) analysis of RNAseq data from indicated cell populations FACS sorted from WT (FOXO1^flox/flox^ FOXO3^flox/flox^) mice (n = 3 per population). The variation explained by each PC is displayed in parenthesis. (C) Venn diagram showing the overlap between expressed protein-coding genes in indicated populations. Genes with ≥0.3 transcript per million (TPM) in all three replicas were considered expressed. (D) Hierarchically clustered heatmaps showing expression of protein-coding genes important for NK cell or ILC development. (E) Volcano plots showing differentially expressed genes for the comparisons between ILCP versus CLP (left panel) and NKP versus ILCP (right panel). Differentially expressed genes (regulated by ≥2-fold at an FDR<0.05) are highlighted in color. Circle sizes indicate expression values (log2(TPM+1). (F) Hierarchically clustered heatmap showing row normalized expression of differential expressed genes (identified in E). Clusters I-IV are indicated. (G) KEGG pathway analysis of differentially expressed genes comparing ILCP versus CLP (left) and NKP versus ILCP (right). Genes regulated by ≥2-fold at an FDR<0.05 were considered differentially expressed and used in the analysis. The size and color of the circles indicate the number of genes in each category and significance of enrichment respectively. (H-I) Expression levels of FOXO1 and FOXO3 from indicated progenitor populations, obtained by (H) RNA-seq or (I) flow cytometry. Dots represent individual analysed animals (*n* = 2-4). Bars indicate mean and SD. Data shown in (I) is from one representative experiment out of two independent experiments.

Next, we investigated the expression pattern of genes known to be crucial for NK and ILC development (Suzuki, Duncan et al. 1997, Zhong and Zhu 2017) **(Fig. 1D)**. A large number of these genes, including *Bcl11b, Tox2, Zbtb16, Tcf7, Rorc, Id2 and IL2rb*, were found to be upregulated at the ILCP stage **(Fig. 1D)**. Looking directly at genes with significantly changes expression in the CLP to ILCP **(Fig. 1E left)** and ILCP to NKP **(Fig. 1E right)** transitions, revealed an overall pattern where ILCPs displayed down-regulation of genes expressed at the CLP stage (**Fig. 1F, cluster II and in part I**) and up-regulate genes expressed NKP stage **(Fig. 1F, cluster III and in part IV)**. When annotated (using Metascape), we as expected found that cluster I and II were enriched for B cell lineage associated genes (Inlay, Bhattacharya et al. 2009, Mansson, Zandi et al. 2010). In contrast, genes in cluster III and IV were enriched for genes associated with the NK cell lineage (Fig. S1A). Hence, suggesting that the B-lineage associated gene program observed in CLPs (Inlay, Bhattacharya et al. 2009, Mansson, Zandi et al. 2010) is shut down in ILCPs. ILCPs instead adopt a general ILC gene program before a more refined NK gene program is established in the NK-lineage committed NKPs (Fathman, Bhattacharya et al. 2011).

To characterize changes in cell-signalling pathways occurring at the developmental transitions, we performed KEGG pathway analysis (**Fig. 1G**) on the differentially expressed genes (**Fig. 1E-F**). This revealed a significant enrichment of genes involved in cytokine-cytokine receptor interaction as well as the PI3K-Akt, MAPK and Rap1 signaling pathways **(Fig. 1G)**. This is in line with prior observations of the critical involvement of cytokines and downstream signaling for early NK cell development and maturation (Ali, Nandagopal et al. 2015, Yang, Li et al. 2015, Wu, Tian et al. 2017, Gotthardt, Trifinopoulos et al. 2019).

Interestingly, the cytokine, PI3K-Akt, MAPK, and Rap1 pathways all coalesced on the FOXO-family. This by either modulating FOXO localization and activity or by altering expression of genes that are direct transcriptional targets of the FOXO-family (Asada, Daitoku et al. 2007, Zhang, Tang et al. 2011, Webb, Kundaje et al. 2016, Birnbaum, Wu et al. 2019). We found that FOXO1 and 3 were co-expressed in Ly6D^neg^ CLP, ILCP and NKP on the mRNA level **(Fig. 1H)**. To further validate this observation, we confirmed that FOXO1 and FOXO3 were expressed at the protein level at all three progenitor stags (Fig. 1I) as well as in committed NK cells from spleen and BM (Fig. S1B-C). This prompted us to further explore of the role of the FOXO family in NK cell development (Deng, Kerdiles et al. 2015, Huang, Wang et al. 2019).

### NK cell development and maturation are dependent on the FOXO proteins

To understand the role of FOXO1 and FOXO3 in NK cell development, we utilized Vav^−^iCre (de Boer, Williams et al. 2003) to conditionally ablate FOXO1 (FOXO1Δ^Vav^) (Paik, Kollipara et al. 2007) and FOXO3 (FOXO3Δ^Vav^) (Castrillon, Miao et al. 2003) individually or in combination (FOXO1,3Δ^Vav^) throughout the hematopoietic system. Littermates lacking Vav-iCre (mainly FOXO1^flox/flox^FOXO3^flox/flox^ animals) were utilized as controls (WT). While residual FOXO1 and FOXO3 proteins were detected in the conditional mice (**Fig. S2A**), the floxed DNA binding domains of both genes were found to be very efficiently deleted by Vav-iCre when investigated at the mRNA level **(Fig. S2B)**.

Neither the loss of FOXO1 or FOXO3 alone resulted in a significant alteration to the number of NK cells in spleen and BM **(Fig. 2A-B)**. In sharp contrast, NK cell numbers were severely reduced in the spleen (5-10 fold reduced) and clearly decreased in the BM of FOXO1,3Δ^Vav^ animals (2-5 fold reduced) **(Fig. 2A-B)**. Altogether, this implies that FOXO1 and FOXO3 display functional redundancy in NK cell development and together, are critical for generation or maintenance of NK cell numbers.

**Fig. 2.**
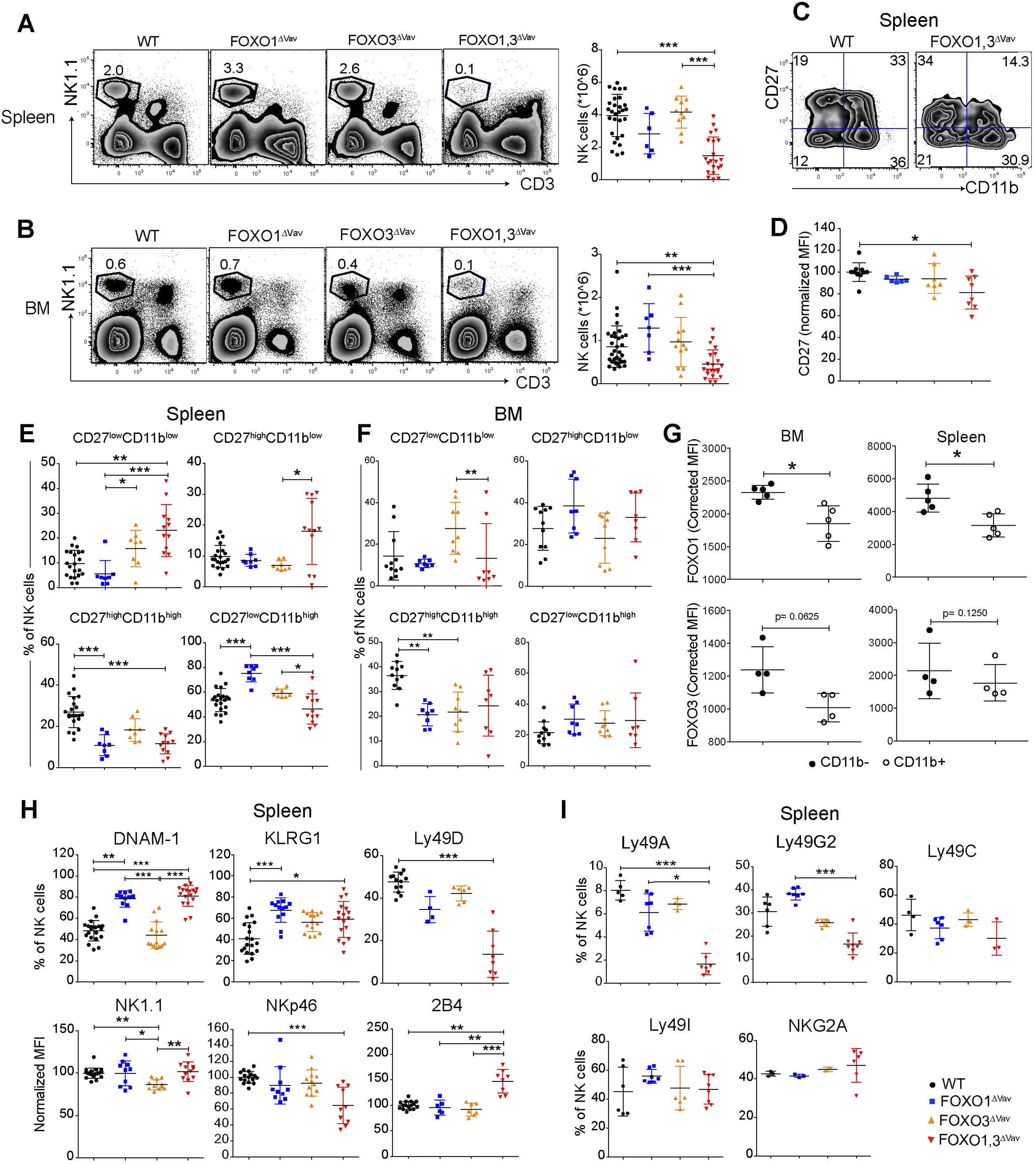
NK cell development is dependent on FOXO1 and FOXO3. (A) Representative flow cytometry profiles (left panel) and total number of NK cells (single, live NK1.1^+^CD3^−^ cells) (right panel) in spleen from animals with the indicated genotypes (n = 6-14). (B) Representative flow cytometry profiles (left panel) and total number of NK cells (single, live NK1.1^+^CD3^−^ cells) in bone marrow (BM) (right panel) from animals with the indicated genotypes (n=7-17). (C) Representative flow cytometry profiles showing splenic NK cell maturation stages in animals with the indicated genotypes. (D) CD27 MFI of CD27^+^ NK cells from spleens (n = 6-12). (E) Frequency (%) of splenic NK cells from each indicated maturation stage (*n* = 7-17). (F) Frequency (%) of BM NK cells from each indicated maturation stage (*n* = 8-11). (G) FOXO1 and FOXO3 protein expression (MFI) in BM and splenic NK cells (n = 4-5) from C57bl6 mice. Data is from one representative experiment out of two independent experiments. (H) Frequency (%) or normalized MFI of indicated activating receptors on splenic NK cells (*n* = 14-19). (I) Frequency (%) of splenic NK cells with indicated inhibitory receptors (*n* = 3-8). In panels A-B and D-I: dots represent individual analyzed animals; bars indicate mean and SD; *, ** and *** indicates p-values <0.05, <0.01 and <0.001 respectively. P-values were calculated using: Kruskal Wallis tests with Dunn’s multiple comparisons test (panels panels A-B, D-F and H-I) or the paired non-parametric Wilcoxon T test (panel G). Symbols utilized to indicate the genotype of analyzed mice throughout the panels are shown in the bottom right corner of the figure.

We next assessed NK cell maturation in the three FOXO-deficient mouse strains according to CD11b and CD27 expression (Hayakawa and Smyth 2006, Luu, Wagner et al. 2019). In this scheme, the first stage of NK cell maturation is characterized by low expression of both CD27 and CD11b (CD27^low^CD11b^low^). CD27 is then increased (CD27^high^CD11b^low^) before the subsequent upregulation of CD11b (CD27^high^CD11b^high^) and finally CD27 being down-regulated again (CD27^low^CD11b^high^) at the last step of maturation. The expression of CD27 on CD27^high^ NK cells was significantly reduced in FOXO1,3Δ^Vav^ as compared to WT **(Fig. 2C-D)** but regardless, all four maturations stages could be distinguished **(Fig. 2C)**. Analysis of the maturation subsets in spleen revealed that FOXO1,3Δ^Vav^ NK cells were less mature compared to NK cells from WT and single knockouts, with an accumulation of the CD27^low^CD11b^low^ population and a significant increase of the CD27^high^CD11b^low^ population **(Fig. 2E)**. Interestingly, there was an accumulation of the terminally differentiated mature CD27^low^CD11b^high^ NK cells in FOXO1Δ^Vav^ mice, that was not observed in FOXO3Δ^Vav^ and FOXO1,3Δ^Vav^ mice **(Fig. 2E)**. This suggests that in the periphery FOXO1 might act as a brake for FOXO3-driven maturation as its absence only has an effect when FOXO3 is present. In contrast to what was found in the spleen, we observed no accumulation of the immature CD27^low^CD11b^low^ subset and only a trend towards mature CD27^high^CD11b^high^ NK cells being reduced in the BM of FOXO1,3Δ^Vav^ mice **(Fig. 2F)**.

To investigate whether the observed phenotype had any relation to FOXO expression, we quantified FOXO protein expression in early (CD11b^low^) and late (CD11b^high^) NK maturation **(Fig. 2G)**. In line with the reduced generation of later CD11b^high^ NK cells in FOXO1,3Δ^Vav^ mice, we found that both FOXO1 and FOXO3 generally displayed higher expressed in the more immature CD11b^low^ NK fraction **(Fig. 2G)**. Together this suggesting that the FOXO proteins to a higher extent influence developmental progression of CD11b^low^ NK cells and that peripheral NK cell development is more dependent on the FOXO proteins then BM NK cell development. The later, in line with the higher FOXO protein expression observed in splenic NK cells (**Fig. S1B-C**).

### Loss of FOXO results in perturbed NK receptor expression

We next investigated the impact of FOXO towards balancing activating and inhibitory receptors that control signaling and functionality in NK cells (Kadri, Wagner et al. 2016, Ganesan, Luu et al. 2017). Loss of FOXO3 caused no significant changes in the NK receptor repertoire with the exception of a reduction in NK1.1 expression **(Fig. 2H-I and Fig. S3A-B)**. In contrast, the loss of FOXO1 alone was enough to cause significant changes in DNAM-1 and KLRG1 expression **(Fig. 2H)**. As FOXO1,3Δ^Vav^ mice displayed a similar phenotype as FOXO1Δ^Vav^ mice **(Fig. 2H)**, this suggests that the perturbation of DNAM-1 and KLRG1 is caused solely by the loss of FOXO1. In addition, FOXO1,3Δ^Vav^ mice uniquely displayed significant changes in the expression of the activating receptors Ly49D, NKp46 and 2B4 **(Fig. 2H and Fig. S4A)** as well as the inhibitory receptors Ly49A and Ly49G2 **(Fig. 2I and Fig. S4B)**. Similar alterations in the activating and the inhibitory receptors were also observed in BM NK cells (data not shown) despite the subtle changes in their maturation status in the mutant mice **(Fig. 2F**). This suggests that the effect of FOXO in the regulation of NK cell repertoire in the spleen might not be related to the maturation status. Hence, the FOXO transcription factors individually or cooperatively influence the expression of specific activating and inhibitory receptors.

### CD122 expression on NK cells is reduced in the absence of FOXO1 and FOXO3

As IL-15 is an important cytokine for NK cell development and survival (Diefenbach, Colonna et al. 2014), we investigated if the reduced expression of CD122, a central component of the IL-15 receptor on NK cells (Suzuki, Duncan et al. 1997) could be a contributing factor to the observed reduction in NK cell numbers in FOXO1,3Δ^Vav^ mice. We found a significant reduction in CD122 expression on both BM and splenic NK cells from FOXO1,3Δ^Vav^ mice **(Fig. 3A)** and that there was a clear correlation between NK cell numbers and CD122 expression in peripheral NK cells **(Fig. 3B)**. Interestingly, dependence on the FOXO proteins differed between BM and splenic NK cells with the CD122 expression on BM NK cells seemingly only relying on FOXO3 **(Fig. 3A)**. In contrast the combined deletion was required to affect CD122 expression in splenic NK cells (**Fig. 3A**), potentially due to the higher expression of FOXO proteins in splenic NK cells as compared to BM NK cells **(Fig. S1B-C)**.

**Fig. 3.**
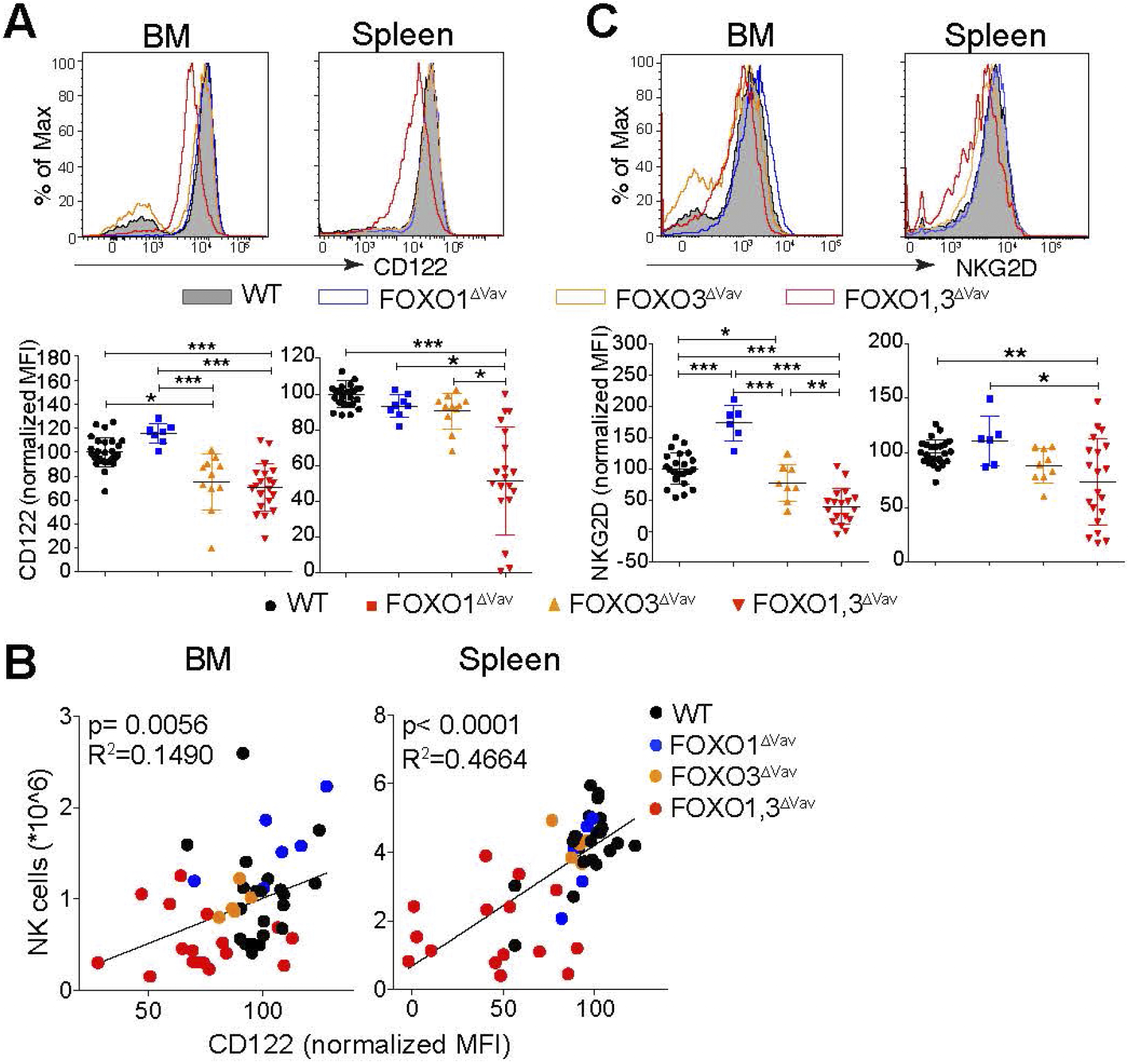
FOXO regulates CD122 expression on NK. (A) Representative flow cytometry profiles (top) and normalized MFI (bottom panel) showing CD122 expression on BM and splenic NK cells (n = 8-28). (B) Correlation between CD122 expression and NK cell numbers (n = 50). (C) Representative flow cytometry profiles (top) and normalized MFI (bottom panel) showing NKG2D expression on BM and splenic NK cells (n = 6-26). In panels A-C: dots represent individual analyzed animals; bars indicate mean and SD; *, ** and *** indicates p-values <0.05, <0.01 and <0.001 respectively. P-values were calculated using: Kruskal Wallis tests with Dunn’s multiple comparisons test (panels A, B) and linear regression (panel C).

It has previously been shown that the expression of the activating receptor NKG2D on NK cells is dependent on IL-15 signaling (Roberts, Lee et al. 2001, Horng, Bezbradica et al. 2007, Luu, Ganesan et al. 2016) and hence it can be utilized as a surrogate marker for IL-15 signaling. Much resembling the expression pattern of CD122 (**Fig. 3A**), we found that NKG2D expression was significantly reduced on both BM and splenic NK ells from FOXO1,3Δ^Vav^ mice but that the dependence on the FOXO proteins varied between BM and spleen (**Fig. 3C**). Hence, we concluded that the loss of FOXO results in reduced surface expression of CD122 and decreased IL-15 signaling. This strongly suggesting, that the reduced NK cell numbers in part can be directly contributed to a diminished IL-15 response in FOXO1,3Δ^Vav^ NK cells.

### FOXO deficiency results in a developmental block at the ILCP to NKP transition

The significant decrease in NK cell numbers (**Fig. 2B**) coupled with the mild phenotype in BM NK cell maturation (**Fig. 2F**) in FOXO1,3Δ^Vav^ mice, hinted at a defect in NK cell progenitors. To investigate this, we performed phenotypic analysis of the CLP, ILCP and NKP compartments **(Fig. 4A)**. This showed a significant decrease in the numbers of CLPs in FOXO1,3Δ^Vav^ mice but no overt changes to the number of ILCPs **(Fig. 4A-B)**. In contrast the downstream CD122^+^ (IL15Rβ) NKP population was significantly decreased in the FOXO1,3Δ^Vav^ mice (**Fig. 4A-B**). Hence, depletion of FOXO results in seemingly increased generation of ILCPs from CLPs and developmental block at the ILCP to NKP transition. The later arguing for that a significant part of the observed reduction in NK cell numbers – in particular in BM - being due to decreased generation of early NK cell progenitors.

**Fig. 4.**
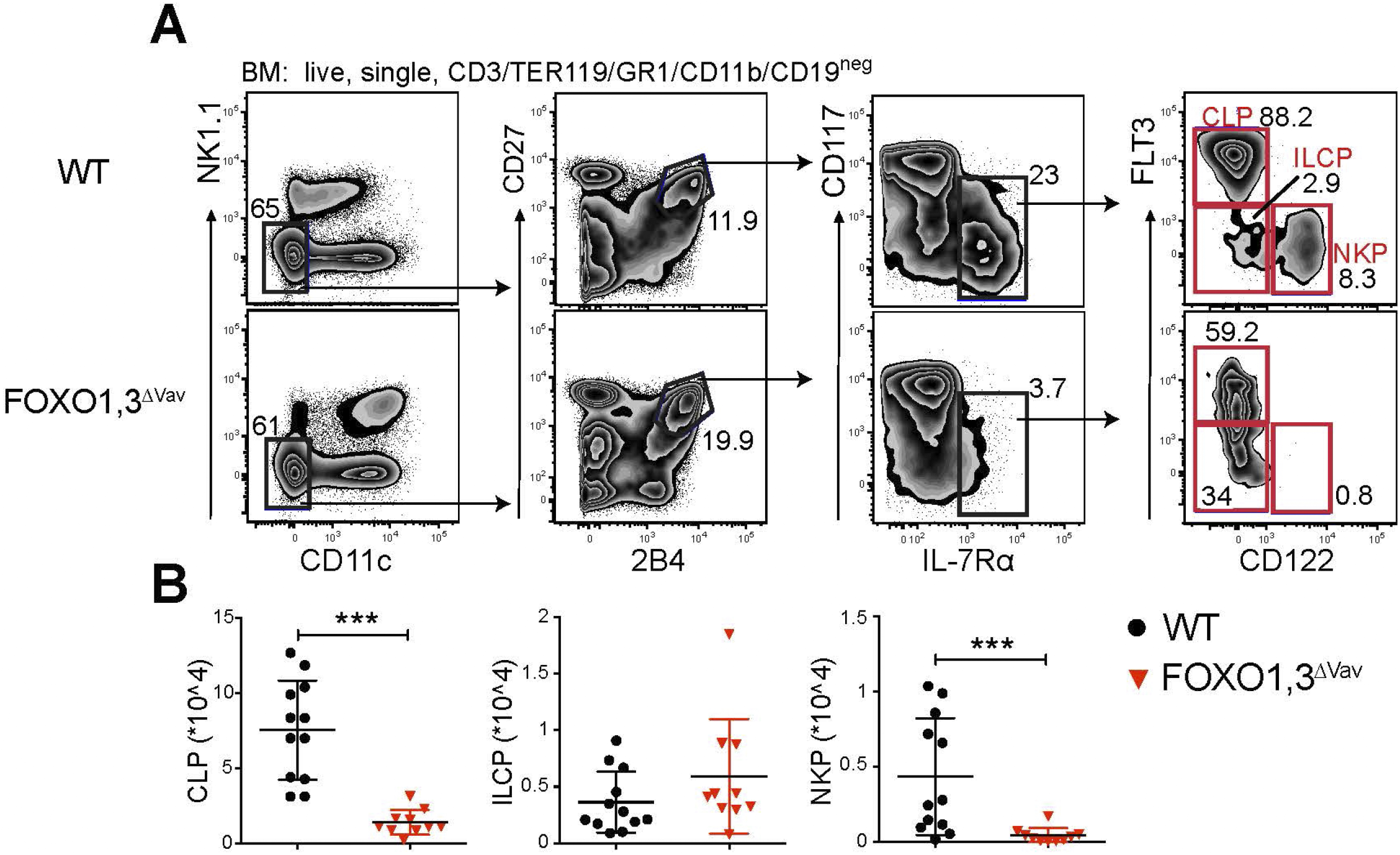
FOXO deficiency results in a block at the ILCP to NKP transition. (A) Representative flow cytometry profiles showing the identification of CLP, ILCP and NKP in animals with the indicated genotype. (B) Total number of CLP, ILCP and NKP in BM of animals with the indicated genotype (*n* = 10-12). In panel B: dots represent individual analyzed animals; p-values were calculated using Mann-Whitney; bars indicate mean and SD; *, ** and ** indicate p-values <0.05, <0.01 and <0.001 respectively.

### The perturbation in NK development is intrinsic to hematopoiesis

To verify that the observed phenotype was intrinsic to hematopoiesis, we utilized the CD45.1/2 system and adoptively transfer FOXO1,3Δ^Vav^ BM (CD45.2) to irradiated WT (CD45.1) hosts. The number of NK cells generated 12 weeks post transplantation from adoptively transferred FOXO1,3Δ^Vav^ BM cells were markedly reduced in spleen **(Fig. S4A and B left)**, BM **(Fig. S4B middle)** and blood (**Fig. S4B right**). The perturbation in NK maturation was also recapitulated with reconstituted FOXO1,3Δ^Vav^ NK cells displaying accumulation of immature (CD27^low^CD11b^low^) NK cells and reduced number of more mature (CD27^high^MAC1^high^) NK cells (**Fig S4C**). Further, we observed a significant reduction in CLP numbers and block at the ILCP to NKP transition in the progeny of FOXO1,3Δ^Vav^ BM cells **(Fig. S4D-E)**. Hence, the presence of normal cells in the transplantation setting do not rescue NK cell development from FOXO1,3Δ^Vav^ donor BM cells, supporting the notion of a cell autonomous FOXO requirement in the regulation of early NK progenitors and NK cell maturation.

### Loss of FOXO impacts expression of NK associated gene already at the CLP stage

With the loss of FOXO perturbing the NK developmental pathway already at the level of the CLP (**Fig. 4A-B**), we next sought to investigate if we could identify NK lineage related changes already at this step of development. To this end, we performed RNAseq on Ly6D^neg^ CLP (GR1/MAC1/NK1.1/TER119/CD3ε^−^CD11C^−^LY6C^−^IL-7Rα^+^FLT3^+^KIT^low^Ly6D^−^) from WT, FOXO1Δ^Vav^ mice, FOXO3Δ^Vav^ mice and FOXO1,3Δ^Vav^ mice. We found 469 genes that displayed significant differential expression between Ly6D^neg^ CLPs from FOXO1,3Δ^Vav^ and WT mice **(Fig. 4A-B and Supplementary Table 2)**. The observed perturbation in gene expression became progressively more distinct with the loss of both FOXO1 and FOXO3 function **(Fig. 4A)**. This suggesting that the FOXO proteins have a mainly synergistic functions at this step of development.

In agreement with earlier rapports, we found that *Il7ra* gene was significantly down-regulated **(Fig. 4A-C)** confirming the known positive regulatory role of FOXO in controlling IL-7R*a* expression (Dengler, Baracho et al. 2008, Ouyang, Beckett et al. 2009). Further, looking specifically at genes previously described to be NK cell signature genes (Goh and Huntington 2017), we found that a significant number of these genes displayed altered expression at the LY6D^neg^ CLP stage **(Fig. 4B-C).** Of note, we found that *Tcf7, Id2, Il18r1*, *Il12rβ1, Cxcl10* and *Cxcl9* – all genes encoding proteins important for NK cell development, migration, and functions (Chan, Perussia et al. 1991, Okamura, Kashiwamura et al. 1998, Thapa, Welner et al. 2008, Delconte, Shi et al. 2016, Jeevan-Raj, Gehrig et al. 2017) - were up-regulated in the FOXO1,3Δ^Vav^ cells. Conversely, looking at the down-regulated genes, we interestingly found that *Ets1* – a gene know to be important for the development of NK progenitors and NK maturation (Barton, Muthusamy et al. 1998, Ramirez, Chandler et al. 2012) - was significantly down-regulated in CLPs **(Fig. 4A-C)**. The down-regulation of ETS1 could also be confirmed at the protein level in CLPs **(Fig. S5A)**.

### The NK associated gene regulatory network in CLPs is perturbed by the loss of FOXO

We next investigated whether the changes in the expression of known NK-cell-development genes could be related also to changes in the gene regulatory landscape. Using the assay for transposase-accessible chromatin followed by sequencing (ATAC-seq), we identified close to 46 000 open chromatin regions across analyzed Ly6D^neg^ CLP with the vast majority existing both in WT and FOXO1,3Δ^Vav^ cells **(Fig. 5D)**. Overall, most of the identified open chromatin regions were localized in intergenic and intronic region **(Fig. 5E, left)**. Next, we identified regions with significant differences (adjusted p-value < 0.01 and ≥2-fold change in signal amongst peaks identified in ≥2 samples and having >30 reads) in chromatin accessibility between WT and FOXO1,3Δ^Vav^. This revealed 297 regions that were mainly localized in intergenic and intronic regions **(Fig. 5E, right)**. Hence, this suggests that loss of FOXO in LY6D^neg^ CLPs results in relatively few but distinct changes to overall chromatin accessibility and that the major effect on gene regulation is via distal elements while promoters remain largely unaffected.

**Fig. 5.**
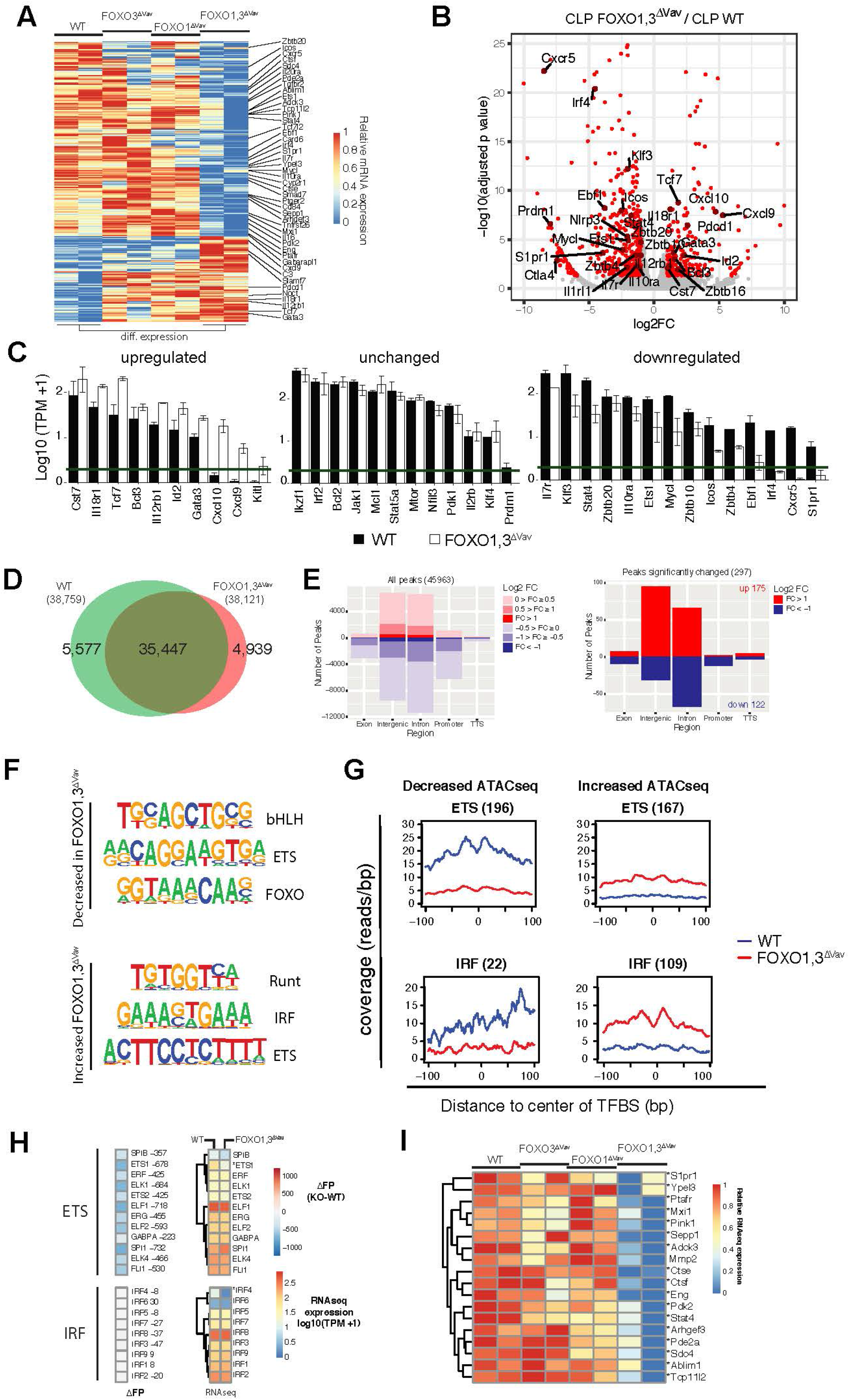
Removal of FOXO results in NK associated gene regulatory changes at the CLP stage. (A) Heatmap showing row normalized expression of genes with differential expression (adjusted p-value ≤0.01, ≥2-fold change in expression and ≥1 TPM in 2+ samples) between WT and FOXO1,3Δ^Vav^ Ly6D^neg^ CLPs. (B) Volcano plot showing log2 fold change and adjusted p-value for the comparison of WT to FOXO1,3Δ^Vav^ Ly6D^neg^ CLPs. Red dots indicate genes with >2-fold change in expression and adjusted p-value < 0.05. (C) Bar graphs showing expression (TPM) of select NK associated genes. Genes with adjusted p-value ≤ 0.01, ≥2-fold change in expression and ≥1 TPM in 2+ samples were considered to have the decreased or increased expression. The green line indicates 1 TPM. (D) Venn diagram showing the overlap between ATACseq peaks identified in Ly6D^neg^ CLPs from WT and FOXO1,3Δ^Vav^ mice. Only peaks identified in ≥2 replicas each with >30 reads were considered. (E) Annotation of all ATAC-seq peaks identified (left) and peaks with significantly altered chromatin accessibility (adjusted p-value < 0.01 and ≥2-fold change in signal) when comparing Ly6D^neg^ CLPs from WT and FOXO1,3Δ^Vav^ mice (right). Number of regions and log2 fold change (FOXO1,3Δ^Vav^ / WT) in ATAC-seq signal is indicated. (F) Motifs enriched in differential ATAC-seq peaks. Top three most significantly enriched motifs existing in >10% of regions are displayed. (G) Cut-profiles for differential ATAC-seq peaks with ETS- and IRF-family transcription factor binding sites (TFBS). Number of regions with each TFBS is indicated in parenthesis. (H) Genome-wide difference in the number of footprints (identified in WT and FOXO1,3Δ^Vav^ LY6D^neg^ CLPs) (left) and expression (right) of indicated genes from the ETS- and IRF-families. * indicates significant differences in gene expression between Ly6D^neg^ CLPs from WT and FOXO1,3Δ^Vav^ mice. (I) Expression of known ETS1 targets in Ly6D^neg^ CLPs from mice with indicated genotypes.

To identify transcription factors whose altered binding could cause the alterations in chromatin accessibility, we performed *de novo* motif enrichment analysis on the peaks with altered chromatin accessibility. This revealed that peaks with decreased accessibility in the FOXO1,3Δ^Vav^ Ly6D^neg^ CLP were enriched for transcription factor binding (TFBS) sites related to the bHLH-, ETS- and, as expected, the FOXO-family **(Fig. 5F top)**. Correspondingly, RUNT, IRF and ETS motifs were found in regions which gained chromatin accessibility **(Fig. 5F bottom)**. To further corroborate that TF binding was altered, we analyzed the Tn5 integration sites (cut-profiles) around the putative TFBS for each of these TF-families. Out of the motifs identified **(Fig. 5F)**, ETS and IRF produced clear cut-profiles **(Fig. 5G and Fig. S5B).** This supporting that altered ETS and IRF binding directly contribute to the changes in chromatin accessibility while suggesting that the other TFBS are either not used or that TF binding does not produce distinct cut-profiles on these analyzed regions.

Altered binding of a single transcription factor might not cause significant changes to the overall chromatin accessibility at the level of a whole chromatin region as defined by the ATACseq peaks. A complementary approach is footprinting analysis, which instead attempts to localize sudden decrease in the number of reads within an open chromatin region to identify individual TF bound regions (Vierstra and Stamatoyannopoulos 2016). By means of footprint analysis, we assessed the changes in genome-wide binding of ETS and IRF. We found no major changes in overall number of IRF associated footprints (**Fig. 5H**). This suggesting that increased IRF binding is specifically associated with peaks displaying increased chromatin accessibility in FOXO1,3Δ^Vav^ CLPs (**Fig. 5F-G**), while the observed decrease Irf4 expression (**Fig. 5C and H**) has no major impact on overall IRF-binding.

In contrast, we found a significant decrease in footprints associated with ETS motifs also at the genome-wide level **(Fig. 5H)**. With Ets1 being the only identified ETS-family member displaying significant changes in expression **(Fig. 5C and H)** this suggests that the decreased number of ETS-bound regions reflects reduced binding of ETS1. This conclusion is further supported by putative ETS1 target genes (Consortium 2004, Consortium 2011) including *Stat4, Pdk2, Adck3* showing significantly lower expression in FOXO1,3Δ^Vav^ CLPs **(Fig. 5I)**. Taken together with the reduced expression of ETS1 (**Fig. 5B-C and Fig. S5A**) and clear ETS cut-profile (**Fig. 5G**), this suggests that loss of ETS1 binding in FOXO1,3Δ^Vav^ CLPs contribute to the altered gene regulatory landscape and potentially the reduced capacity to generate NK cells downstream of the CLP (Ramirez, Chandler et al. 2012).

We further looked specifically at TCF7 (Jeevan-Raj, Gehrig et al. 2017) and NFIL3 (Male, Nisoli et al. 2014) as both are known to be critical for BM NK progenitor development. TCF7 expression was increased in CLPs from FOXO1,3Δ^Vav^ mice **(Fig. 5B-C)** but no significant change in binding as assessed by footprinting analysis (46 less footprints in the FOXO1,3Δ^Vav^ CLPs) was observed. Hence, TCF7 is potentially controlled by FOXO at the transcriptional level but only have minor impact in terms of chromatin accessibility at the CLP stage. NFIL3 showed no significant change in expression or overall binding (7 less footprints in the FOXO1,3Δ^Vav^ CLPs). Hence, we find no indication that perturbed TCF7 and NFIL3 activity contribute to the changes observed in FOXO1,3Δ^Vav^ CLPs.

### ILCPs lacking FOXO fails to up-regulate NK-lineage related genes

With FOXO1,3Δ^Vav^ mice displaying a block at the ILCP to NKP transition, we next sought to characterize the transcriptional changes caused by the loss of FOXO in ILCPs. Based on PCA, the FOXO1,3Δ^Vav^ ILCPs overall maintained an ILCP transcriptional profile as compared to its WT counterparts (PC2) (**Fig. 6A**). In line with this, the expression of Id2 – which marks the formation of ILCP from CLP (Xu, Cherrier et al. 2019) – was not altered and generally the expression of ILC related transcription factors (Zhong and Zhu 2017) was also found to be similar **(Fig. 6B)**. However, the combined loss of FOXO1 and FOXO3 did cause distinct gene expression changes as observed both by PCA (PC1) (**Fig. 6A**) and direct comparison of expression profiles (**Fig. 6C and supplementary table 1**). In relation to normal development, the FOXO1,3Δ^Vav^ ILCPs displayed lower expression of genes commonly expressed throughout the early NK progenitor hierarchy (**Fig. 6D, cluster I**) and to a lesser extent maintained expression of CLP associated genes (**Fig. 6D, cluster IV**). We in addition found that the FOXO1,3Δ^Vav^ ILCPs failed to properly express genes normally upregulated in the CLP-ILCP transition and then further increased in expression in the ILCP-NKP transition (**Fig. 6D, cluster II**). The later gene cluster was as expected enriched for NK lineage associated genes (**Fig. S6A**) including amongst others *Klrd1* (encoding for CD94) and *Tigit* shown to function as NK immune checkpoint inhibitors (Khan, Arooj et al. 2020). The reduced generation of NKPs from FOXO1,3Δ^Vav^ ILCPs is hence associated with a failure to properly express an early NK gene program.

**Fig. 6.**
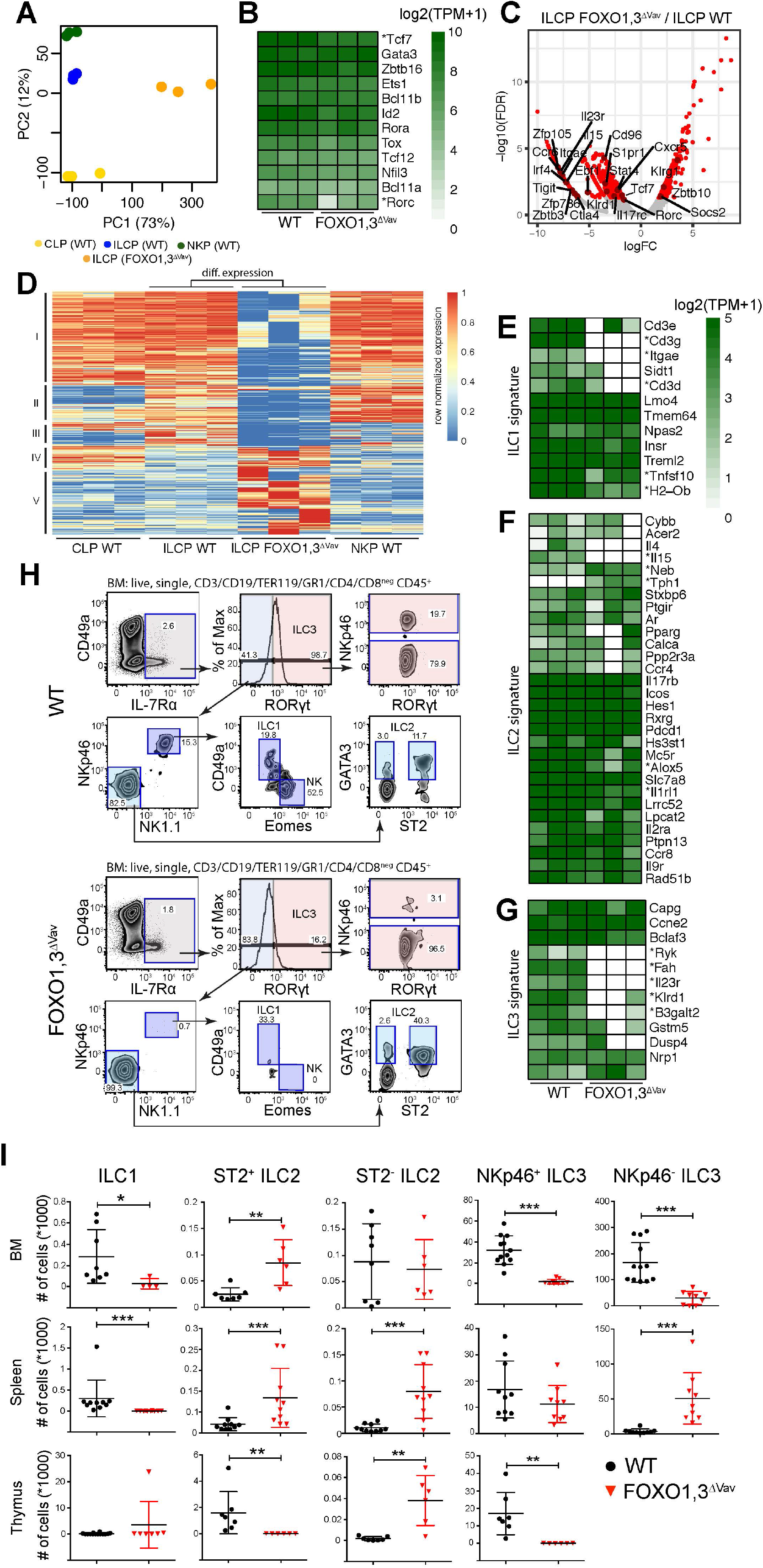
Loss of FOXO results in a perturbation of the ILCP transcriptional program and the development of non-cytotoxic ILCs. (A) Principal component (PC) analysis of RNAseq data from indicated cell populations (for gating strategy see Fig. 1A.) from wildtype and FOXO1,3Δ^Vav^ mice (n = 3). The variation explained by each PC is displayed in parenthesis. (B) Hierarchically clustered heatmaps showing gene expression for selected transcription factors critical for NK/ILC development. * indicates significant differences in expression between WT and FOXO1,3Δ^Vav^ ILCPs (FDR < 0.05, >2-fold change). (C) Volcano plot showing log2 fold change and adjusted p-value for the comparison of WT to FOXO1,3Δ^Vav^ ILCPs. Red dots indicate genes with >2-fold change in expression and adjusted p-value < 0.05. (D) Hierarchically clustered heatmap showing row normalized expression of differential expressed genes (identified in C). Clusters I-V are indicated. (E-G) Hierarchically clustered heatmaps showing gene expression of (E) ILC1, (F) ILC2 and (G) ILC3 signature genes defined by Robinette et al., (Robinette, Fuchs et al. 2015). * indicates significant differences in expression between WT and FOXO1,3Δ^Vav^ ILCP (FDR < 0.05, >2-fold change). (H) Representative flow cytometry profiles showing the identification of ILC subsets in BM from WT and FOXO1,3Δ^Vav^ mice. (I) Total number of indicated ILC subsets in BM, spleen and thymus from WT and FOXO1,3Δ^Vav^ mice (n = 6-12). NKp46-ILC3 population had less than 50 cells per thymus in all mice from both mouse strains so was not shown. Dots represent individual analyzed animals. Bars indicate mean and SD. P-values were calculated using Mann-Whitney tests with *, ** and *** indicates p-values <0.05, <0.01 and <0.001 respectively.

Looking specifically at individual genes impacting NK development, we found that TCF7 - in contrast to what was observed in CLPs (**Fig. 5B-C**) - was downregulated in FOXO1,3Δ^Vav^ ILCPs while NFIL3 expression remained unaffected (**Fig. 6B-C**). Hence, the lower TCF7 expression in ILCPs could potentially contribute to the impaired generation of NKP (Jeevan-Raj, Gehrig et al. 2017). Further, in line with data from the CLP stage, we found that ETS1 displayed a clear trend towards being down-regulated in FOXO1,3Δ^Vav^ ILCPs (**Fig. S6B and 6B**). This in concordance with the CLP expression and epigenetic data arguing for the loss of FOXO causing ETS1 downregulation which in turn contributes to the observed impairment generation of NK cells. Further in line with perturbed ETS1 expression contributing to the observed NK phenotype, we found that Ets1 expression was significantly down-regulated also in splenic NK cells (**Fig. S6C).**

Interestingly, we also found a significant decrease in expression of zink finger protein 105 (*Zfp105)* **(Fig. 5B)**– a transcription factor previously shown to regulated by FOXO1 (Ouyang, Liao et al. 2012, Kim, Ouyang et al. 2013) and to augment differentiation towards NK cell lineage (Chambers, Boles et al. 2007) as well as being a direct target of FOXO1 in CD8 T cells – which might suggest that FOXO-mediated regulation of Zfp105 expression plays a role in NK cell development.

### Loss of FOXO1 and FOXO3 disrupts development of non-cytotoxic ILC

ILCP represents a heterogenous population which give rise not only to NK cells but also to the other non-cytotoxic ILC subsets (Constantinides, McDonald et al. 2014, Xu, Cherrier et al. 2019). In agreement with this, we observed expression of genes encoding transcription factors linked to development of the also non-cytotoxic NK cells, including *Tcf7, Tox, Bcl11b, Zbtb16, Rorc, Ets1, Nfil3* (Zhong and Zhu 2017) in both WT and FOXO1,3Δ^Vav^ ILCP **(Fig. 6B)**.

We next sought to investigate if we could observe transcriptional changes in the FOXO1,3Δ^Vav^ ILCP indicative of disruptions potentially influencing also the non-cytotoxic ILC subsets. To this end, we utilized the ILC1-3 gene signature published by Robinette *et al.* (Robinette, Fuchs et al. 2015). Interestingly, we found down regulations of expression across all the three gene signatures (**Fig. 6E-G**) but most notably within the ILC1 (**Fig. 6E**) and ILC3 (**Fig. 6F**) gene signatures.

These transcriptional changes made us speculate that the development of the non-cytotoxic ILC subsets in addition to NK cells could be perturbed in FOXO1,3Δ^Vav^ mice. To investigate this, we performed phenotypic analysis of ILC subsets from BM, spleen and thymus from WT and FOXO1,3Δ^Vav^ mice (**Fig. 6H**). Indeed, we found that the ILC1 population was reduced in BM and spleen (**Fig. 6H-I**). ILC3 numbers were similarly reduced in BM and thymus while in contrast NKp46^−^ILC3 being increased in spleen (**Fig. 6H-I**). In contrast, ILC2 subsets were generally increased in the analyzed organs (**Fig. 6H-I**). Hence, we concluded that the loss of FOXO causes broad perturbation of the non-cytotoxic ILC subsets. Potentially these changes are due to gene regulatory changes already at the level of the ILCP, meaning that FOXO determine lineage specification of ILC.

## DISCUSSION

In this study, we show that FOXO1 and FOXO3 are expressed in the early progenitors of the innate lymphoid lineages and cooperatively regulate the generation of NK cell progenitors and NK cells. In addition, we discovered a hitherto undescribed role of the FOXO family in establishing the NK/ILC gene expression program in progenitor cells and in the development of the ILC1, 2 and 3 subsets. Hence, the loss of FOXO1 and FOXO3 disrupts development of both the cytotoxic and non-cytotoxic ILC lineages.

Using a combination of RNAseq and ATAC-seq data to study the underlying gene regulatory mechanisms, we found that the loss of FOXO proteins disrupted the regulation of NK and ILC associated genes already at the CLP stage and more markedly so at the ILCP stage. Likely the failure to establish the NK/ILC gene program in ILCPs directly result in the observed reduction in NKP. Interestingly, we found a decrease in ETS1 at the CLP stage onwards in FOXO1,3Δ^Vav^ mice. In line with ETS1 being a critical downstream target of FOXO, the NK cell phenotypes of the ETS1 knockout very much resemble the FOXO1,3Δ^Vav^ phenotype. This, with ETS1 deficient mice displayed reduced splenic NK cells and NKP while ILCPs we seemingly unaffected (Barton, Muthusamy et al. 1998, Ramirez, Chandler et al. 2012). Further, altered NKp46 and Ly49D expression were also observed in ETS1 deficient animals (Ramirez, Chandler et al. 2012). With the activity of ETS1 being modulated via interaction with FOXO1 (Boccitto and Kalb 2011), the loss of FOXO could mimic the ETS1 knock-out by both lowering ETS1 expression and ETS1 activity throughout NK cell development.

The ILCP compartment is heterogeneous and contains several progenitor populations out of which only a subset is involved in the generation of the NK lineage (Constantinides, McDonald et al. 2014, Xu, Cherrier et al. 2019). While these subsets are yet to be readily identifiable without the use of reporter genes, we observed distinct changes both in the activation of the overall ILCP gene expression program and in genes associated with the ILC1-3 lineages. These changes were associated with decreased numbers of ILC1 and ILC3 as well as increased numbers of ILC2. Hence, this argues that the altered gene expression caused by the loss of FOXO is directly reflected in the ILC lineages though it is unclear if these alterations reflect changes at the progenitor composition of the ILCP compartment or overall transcriptional program. Addressing this point will require further studies using reporter mice to distinguish the different progenitor populations within the ILCP compartment.

The loss of FOXO could also directly influence later the development of ILC (i.e : by acting downstream of important ILCP transcription factors). Indeed, the expression of BCL11B in ILCP, especially in combination with ZBTB16, marks the development of ILC2 (Xu, Cherrier et al. 2019). On the other hand, the absence of both markers in ILCP allows for a balanced development of NK cells and all ILC lineages (Xu, Cherrier et al. 2019). FOXO proteins have been suggested to repress cell cycle progression downstream of BCL11B, which is a critical regulator of basal cell quiescence in the mammary gland (Cai, Kalisky et al. 2017). ZBTB16 overexpression leds to reduced FOXO phosphorylation (Chen, Qian et al. 2014) while FOXO1 expression is induced in ZBTB16 heterozygosity as compared to homozygosity (Liska, Landa et al. 2017). Altogether, this makes it tempting to speculate that FOXO serve as crucial transcriptional regulator acting downstream of ZBTB16 and/or BCL11B to control ILC development in a cell type-specific manner.

Prior studies using Ncr1-iCre and VAV-iCre have reported contradictory data on the role of the FOXO1 in NK cell development (Deng, Kerdiles et al. 2015, Wang, Xia et al. 2016). Using the VAV-iCre model, we in agreement with Deng et al., found that the deletion of FOXO1 did not significantly alter total NK cell numbers while CD27^low^CD11b^high^ NK cells were accumulated in line with FOXO1 suppressing Tbx21 expression needed for maturation (Deng, Kerdiles et al. 2015). However, in contrast to what was observed following the combined deletion of FOXO1 and FOXO3 in the Ncr1-Cre model, we found that NK cell numbers declined significantly in both the BM and spleen of FOXO1,3Δ^Vav^ animals. The splenic NK cells that developed displayed an accumulation of immature CD11b^low^ NK cells but lacked the accumulation of CD27^low^CD11b^high^ NK cells observed after the deletion of FOXO1 alone. In addition, the FOXO1,3Δ^Vav^ NK cells displayed perturbed expression of several inhibitory and activating receptors. The discrepancy in maturation status between FOXO1,3Δ^Ncr1^ and FOXO1,3Δ^Vav^ might be attributed to their differential roles in hematopoietic progenitors and committed NK cells.

Somewhat surprisingly, BM NK cell maturation in the FOXO1,3Δ^Vav^ animals was left rather unperturbed. Potentially, this argues for the lower NK cell numbers in BM being a consequence of the reduced numbers of NKPs rather than major issues with NK cell maturation. Speculatively, this would in addition suggest that NK maturation in spleen and BM to some extent have different requirements and that the environment causes different reliance on the FOXO proteins.

IL-15 signalling is critical for NK cell development and survival (Lodolce, Boone et al. 1998, Kennedy, Glaccum et al. 2000, Ranson, Vosshenrich et al. 2003). We found that the FOXO1,3Δ^Vav^ NK cells displayed markedly reduced CD122 (IL15Rβ) and that expression was quantitatively correlated with NK cell numbers. Supporting the notion that the FOXO1,3Δ^Vav^ NK cells have reduced IL-15-signaling, we also found that the IL-15 dependent expression of NKG2D (Roberts, Lee et al. 2001, Horng, Bezbradica et al. 2007, Luu, Ganesan et al. 2016) was significantly lower on BM and splenic NK. Hence, this argues that the reduced NK cell numbers in FOXO1,3Δ^Vav^ mice results from defects in generation of NK cells via NKPs in combination with decreased ability to respond to IL-15 signalling critical for maintaining the normal NK population. Furthermore, the reduction of the ILC1 population in the absence of FOXO1,3 could be explained to occur similarly to the reduction of the NK cell population through their indispensable requirement of IL-15 needed for cell development (Suzuki, Duncan et al. 1997, Vosshenrich, Ranson et al. 2005).

It is appealing to speculate that the changes in progenitor development as well as the reduced CD122 expression can be attributed to the reduction in ETS1 expression. Indeed, chromatin immune precipitation experiments revealed that ETS1 binds to the promoter of the CD122 gene and CD122 expression was reduced in mature ETS1 KO NK cells (Ramirez, Chandler et al. 2012). FOXO1 and FOXO3 hence might tune CD122 expression indirectly through regulating a network of factors including ETS1 to gradually modulate IL-15 responsiveness. rather than causing an “on/off” situation in CD122 expression. Given that once NK cells acquire NKp46 expression (and hence gene deletion occurs in the Ncr1-Cre model), CD122 expression is not perturbed by the loss of FOXO1 and FOXO3 activity (ref: Deng et al), this suggests that the loss of FOXO activity in FOXO1,3Δ^Vav^ NK progenitors or very early NK cells (prior to Ncr1-Cre mediated deletion) cause a defect that cannot be corrected in later stages of NK cell development. This hypothesis is in line with recent studies showing that IL-15 signalling creates a positive regulatory loop to modulate expression of its receptors and several components of the IL-15 signalling pathway (Yang, Li et al. 2015). Hence, FOXO1 and FOXO3 would then serve as crucial early regulator of NK cell fate by establishing proper IL-15 receptor expression.

In conclusion, the co-expression and regulatory function of FOXO1 and FOXO3 is critical throughout the NK cell development and maturation. Mechanistically, we propose that FOXO1 and FOXO3 – amongst other genes - control the expression ETS1 and CD122 that are both integral for NK cell development. In addition, FOXO proteins selectively promote the development ILC1 and ILC3 but not ILC2. The very well controlled intrinsic modes of NK cell development, differentiation, and maturation by FOXO1 and FOXO3 revealed in our study can drive future efforts to develop anti-tumor and anti-viral immunotherapies targeting FOXO proteins.

## MATERIALS AND METHODS

### Mice

To generate animals conditionally lacking FOXO1 and/or FOXO3 throughout the hematopoietic system, we crossed *FOXO1*^*ff*^ (ref Paik Cell 2007) and/or *FOXO3*^*ff*^ (Castrillon, Miao et al. 2003) with Vav^−^iCre (de Boer, Williams et al. 2003) mice. All alleles were maintained on a C57BL/6 background and mice were predominantly analyzed at 8-14 weeks of age. Congenic CD45.1 WT C57BL/6 mice were used as recipients in transplantation experiments. All animal experiments were approved by the local animal ethics committee.

### Flow cytometry

Single-cell suspension of bone marrow, spleen, blood or thymus were incubated with Fc blocker anti-FcγRIII (2.4G2) and subsequently stained with fluorescent antibodies (Supplementary Table 3) and viability markers (LIVE/DEAD® Fixable Aqua Dead Cell Stain Kit or propidium iodide, both from Invitrogen). Stainings were done at 4°C in PBS with 2 % FBS for 20 min. Results were acquired mainly using the BD LSRFortessa™ or BD FACSymphony™ Flow Cytometers (BD Biosciences).

For FACS sorting of progenitor cells, mature cells were depleted using antibodies against TER119, CD3, CD19, GR1 and MAC1 together with sheep anti-rat IgG Dynabeads (Invitrogen) prior to staining with fluorescent antibodies. Cell sorting was performed mainly on a FACSAriaIIu or FACSAria Fusion (BD Biosciences).

Further analysis of FACS data was performed using Flowjo v9.9.6 (TreeStar, Ashland, OR). Corrected mean fluorescence intensities (MFI) were calculated by subtracting the MFI of control sample (stained only with secondary antibody) from the MFI of the FOXO stained sample. Normalized MFIs were calculated by dividing MFIs with the average MFI observed for WT samples in each independent experiment.

### Transplantation assay

To determine in vivo NK lineage output from FOXO1,3Δ^Vav^ hematopoietic stem- and progenitor cells, unfractionated bone marrow (0.2 × 10^6^ WT or 3 × 10^6^ FOXO1,3Δ^Vav^ CD45.2 donor cells), were injected intravenously into irradiated (950cGy) CD45.1 recipients. Number of unfractionated BM cells transplanted were proportional to the frequency of phenotypic hematopoietic stem cells in WT and FOXO1,3Δ^Vav^ mice (data not shown). Part of the FOXO1,3Δ^Vav^ transplanted animals were in addition given 0.2 × 10^6^ unfractionated WT BM cells as support. Reconstitution was analyzed at 12 weeks post-transplantation using flow cytometry.

### TotalScript based RNAseq and analysis

RNA sequencing was done using TotalScript (Epicenter) on RNA prepared from approximately 5000 FACS sorted cells using RNeasy micro (Qiagen) as previously described (Bouderlique, Pena-Perez et al. 2019). Libraries were sequenced pair end (2×50 cycles) on the Illumina platform. Reads were mapped to the mouse reference genome (mm10) using STAR v2.3.2b (https://github.com/alexdobin/STAR). Strand-specific reads in exons was quantified using HOMER and assessment of differential gene expression analysis was done using EdgeR (Robinson, McCarthy et al. 2010) on raw read count. Data visualizations was mainly done using ggplot2, pheatmaps and R base graphics.

### ATAC sequencing and analysis

ATAC sequencing was performed (using 3000-5000 FACS sorted cells) as previously described (Jin, Chen et al. 2018). Libraries were sequenced pair-end (2×50 cycles) on the Illumina platform (Illumina). Reads were trimmed (using Trim Galore v0.4.1), mapped to the mouse reference genome (mm10) (using Bowtie2 v2.3.3.1) and PCR duplicates removed when making HOMER tag directories (using makeTagDirectory with - tbp 1). Peaks were subsequently identified in sub-nucleosomal reads (read-pairs within 100bp) using HOMER’s findPeaks.pl. Peaks with differential chromatin accessibility were identified using EdgeR on raw read counts in identified peaks. Peaks displaying an adjusted p-value ≤ 0.01 and ≥2-fold change in read count were considered to have differential chromatin accessibility. Only peaks identified in ≥2 replicas each with >30 reads were considered in the analysis. Annotation and motif enrichment analysis of differential peaks were done using the HOMER’s annotatePeaks.pl and findMotifsGenome.pl with - size given respectively.

To make cut-profiles, the localization of known HOMER transcription factor binding sites (TFBS) belonging to members of the enriched TF family (identified by the motif enrichment in differential ATACseq peaks) were localized in the genome using HOMER’s findMotifsGenome.pl. To take into account the position of the Tn5 integration into the genome, custom HOMER tag directories were made off-setting reads on the plus and minus strand with +4 and −5 bases respectively (Ref Buenrostro et al, org ATACseq paper). Read depth centered around TFBS from a specific family were subsequently plotted using HOMER’s annotatePeaks.pl with a - fragLength of 9 (corresponding to the bp covered by Tn5) and - hist 1 (1 bp bins).

Genome-wide footprinting to identify TF binding sites was performed using DNase2TF. In brief, Bowtie2 mapped reads were deduplicated (using Picacard tools’ MarkDuplicates), data from the same population/genotype merged (using SamTools’ merge) and down-sampled to 39 million read-pairs per sample (using Picard Tool’s DownsampleSam.jar). Localization of reads were off-set in the .bam file to take into account the Tn5 integration (as described above) using custom scripts. HOMER tag directories and peak finding was done as described above. Peak files and downsampled .bam files were subsequently used as input for DNase2TF (Sung, Guertin et al. 2014). Identified footprints with a p-value ≤ 0.05 were overlapped with the TFBS identified in the mouse reference genome (mm10) using the transfac catalogue as previously described (Neph, Stergachis et al. 2012). Footprints were associated with TFBS when the center of the TFBS fell within the footprint.

Data visualizations was mainly done using ggplot2, pheatmaps and R base graphics.

### RTqPCR

NK cells were FACS sorted using BD AriaIII (BD Biosciences). CD117+ BM cells were enriched using CD117 MicroBeads, mouse (Miltenyi). RNA was purified using RNeasy micro kit (Qiagen) and cDNA prepared using MultiScribe Reverse Transcriptase (Life Technologies) or SuperScript II (Life Technologies) in combination with random hexamer priming. qPCR was performed using TaqMan™ Universal PCR Master Mix (Life Technologies) and TaqMan probes against: FOXO1 (Mm00490672_m1), FOXO3 (Mm00490673_m1), hprt (Mm01545399_m1 or Mm00446968_m1) and Ets1 (Mm01175819_m1).

### SMART-Seq based RNAseq and analysis

200-500 progenitor cells were FACS sorted into lysis solution with DNase I from the Single Cell Lysis Kit (Invitrogen) and samples prepared according to manufacturer’s instructions. RNAseq libraries were subsequently prepared using the SMART-Seq Stranded Kit (Takara) according to the manufacturer’s instruction. Quality of cDNA library was determined using a Agilent Bioanalyzer according to the manufacturer’s protocol. Libraries were quantified by using the KAPA-SYBR FAST qPCR kit (Roche) and sequenced pair-end (2×75 cycles) on the Illumina NextSeq 500.

Adaptor sequences were trimmed and low-quality reads removed using Trimmomatic (v.0.36). All sequencing reads aligning (HiSAT2, v.2.1.0) to annotated mouse ribosomal RNA genes were discarded. High-quality and ribosomal RNA depleted sequencing reads were aligned to the genome GRCm38.p6/mm10 genome using HiSAT2. Using sorted bam files (Samtools v.1.10), the number of aligned reads were counted (featurecount in subread package v. 2.0.0). After normalization (TMM: trimmed mean of M-values), a differential gene expression analysis (edgeR v. 3.28.1) was performed. Significant differentially expressed genes was distinguished by a false discovery rate (FDR) < 0.05. Data were plotted using ggplot2 (v.2.3.3) in R (v. 3.6.1). Gene ontology analysis was conducted using clusterprofiler (v.3.14.3) with the database org.Mm.eg.db (v.3.10.0) in R. All scripts used for processing of SMART-seq data are deposited on Github: https://github.com/jonasns/NK_FOXO.

### Statistics

Statistical analysis and plotting of FACS data was performed using Graphpad Prism version 6 for Mac OSX (Graphpad Software). Statistics and visualization pertaining to RNAseq and ATAQseq data was performed using sequencing using R version 3.3.3 (R Development Core Team 2008) or assessed as described above.

### Data availability

Total Script based RNAseq and ATACseq data available from ENA under accession number PRJEB20316 and PRJEB41018. SMARTseq based RNAseq data is available from _ under accession number _.

## Acknowledgements

We thank the staff of the animal facilities (Karolinska Institute, Huddinge) for animal care. We would like to also thank the MedH Flow Cytometry core facility (Karolinska Institutet) and the Centre for Cellular Analysis (CCA, Karolinska University Hospital), for providing cell analysis and cell sorting services. We also acknowledge the Center for Hematology and Regenerative Medicine (HERM), Karolinska Institute, for giving a great scientific environment. This work was supported by grants from The Swedish Cancer Society, Swedish Research Council, King Gustav V Jubilee Fund, The Karolinska Institutet Foundations, The Stockholm County Council, The Swedish Foundation for Strategic Research, The Knut and Alice Wallenberg Foundation and generously, a donation by Björn and Lena Ulvaeus. In addition, the Karolinska Institutet doctoral education program (KID) supported the doctoral studies of LPP.

## Author contributions

LTT performed experiments, analyzed data and contributed to writing the manuscript JNS performed RNAseq experiments, analyzed RNAseq data and contributed to writing the manuscript. LPP analyzed RNAseq data, analyzed ATACseq data and contributed to writing the manuscript. SK FACS sorted cells and assisted with animal experiments. AK and CG performed RNAseq and ATACseq experiments. SM and LS assisted with flow cytometry staining, in vitro experiments and manuscript discussion. NF, YH and TB assisted with animal experiments. MK provided critical input on the ATAseq analysis. AKW and BJC helped with functional assays and manuscript discussion. AA and CK assisted with RNAseq experiments, bioinformatic analysis and discussion of the manuscript. PH, RM and NK supervised the study. RM and NK designed the study, performed experiments, analyzed data and wrote the manuscript. All authors commented on the final manuscript.

---

**Fig. S1.**
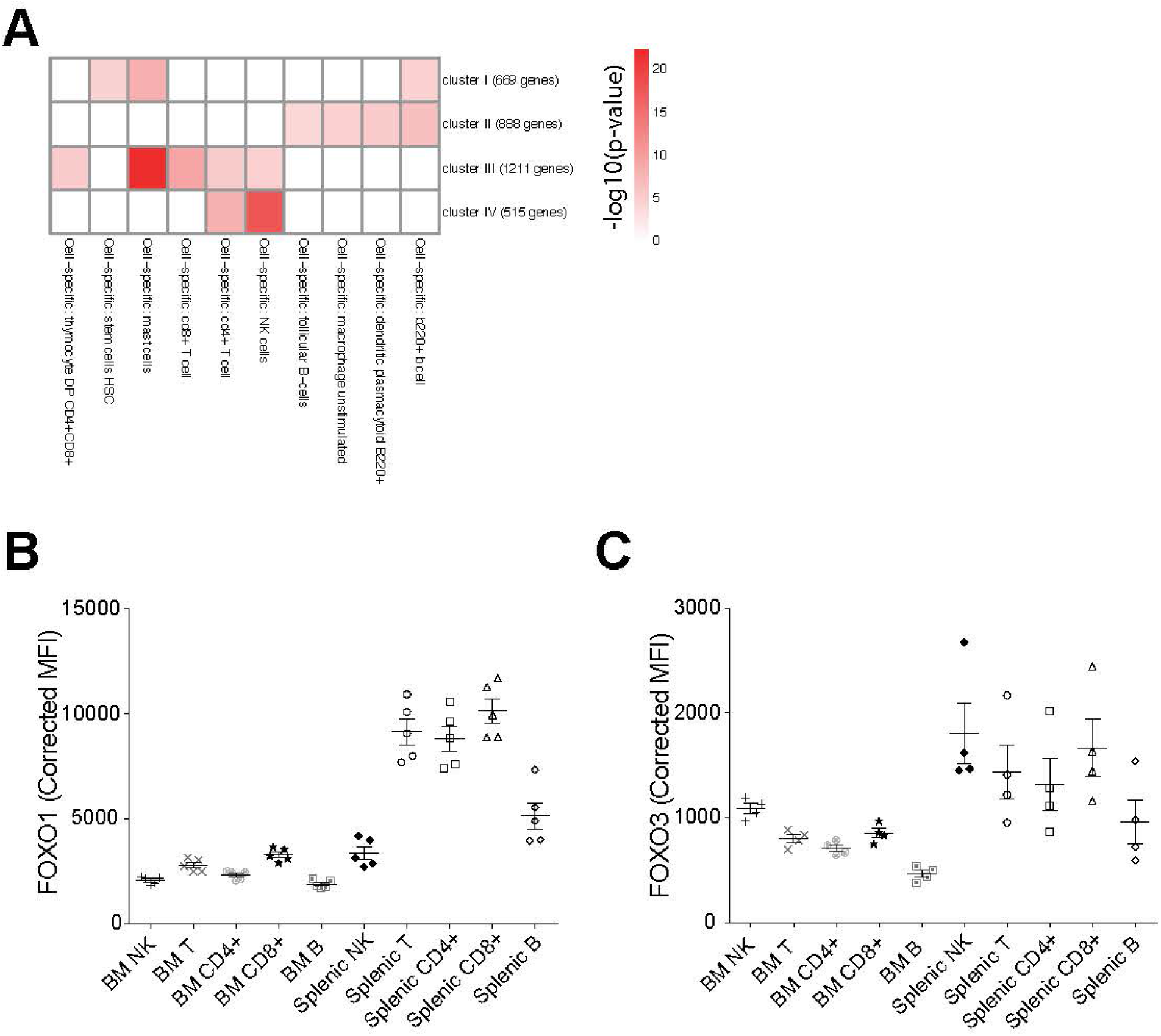
Annotation of developmentally regulated genes and comparison of FOXO expression. (A) Metascape gene enrichment analysis on the gene clusters from Figure 1F. (B-C) FOXO1 (B) and FOXO3 (C) expression in NK cell subsets, T cell subsets, and B cells from C57Bl6 mice as measured by FACS in bone marrow and spleen samples. Dots indicate individual mice. Data is from one representative out of two experiments (*n* = 4-5). Bars indicate mean and SD.

**Fig. S2.**
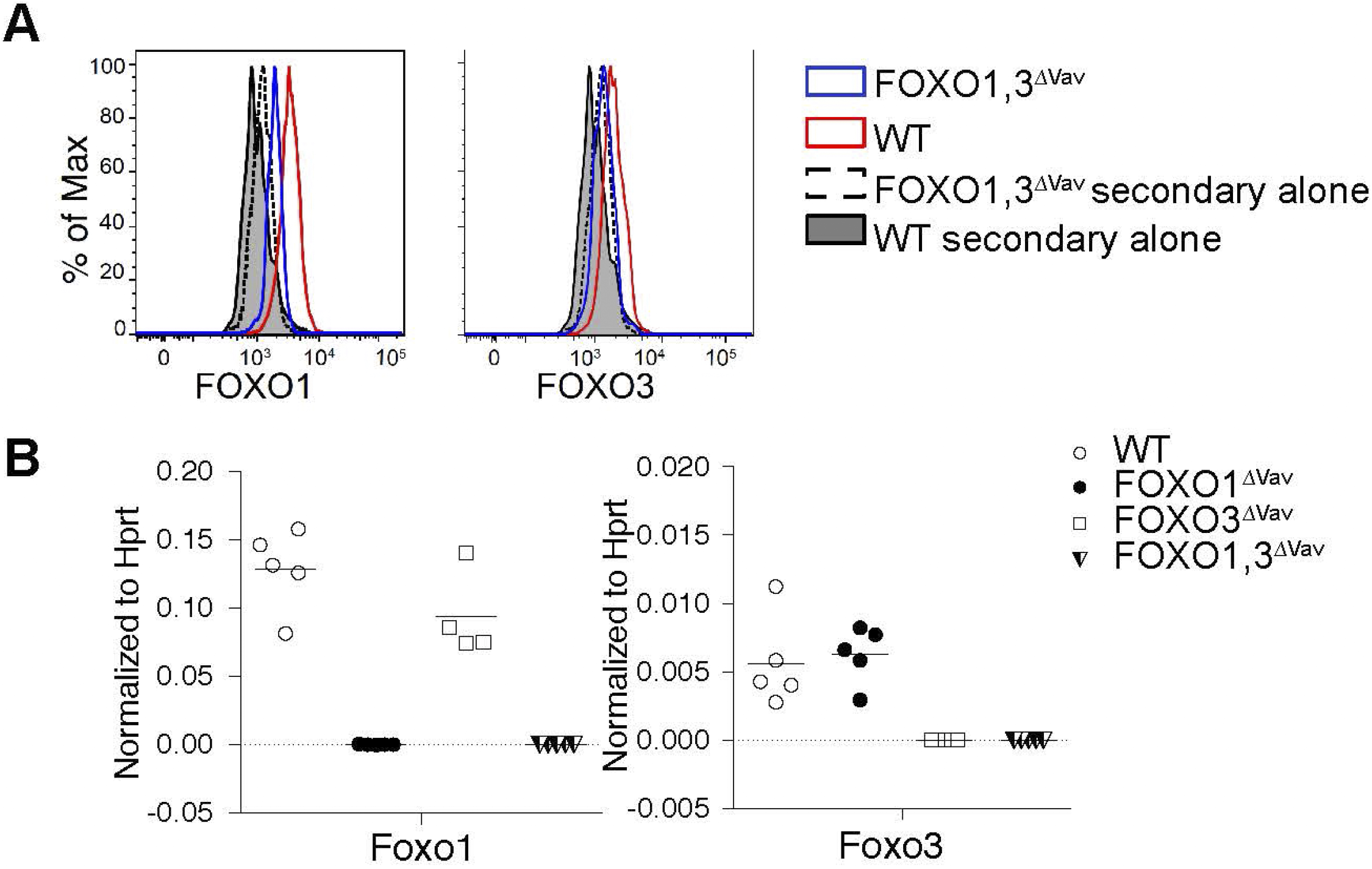
Vav-iCre mediates effective deletion of the loxP flanked FOXO1 and FOXO3 regions. (A) FOXO1 and FOXO3 protein levels in splenic NK cells as measured by intracellular flow cytometry. (B) Expression of FOXO1 and FOXO3 mRNA in mice with the indicated genotype. Each dot represents data from an individual mouse. mRNA levels were measured in KIT (CD117) enriched bone marrow progenitors using RT-qPCR with probes specific to the loxP flanked regions coding for the DNA binding domains of FOXO1 and FOXO3 respectively.

**Fig. S3.**
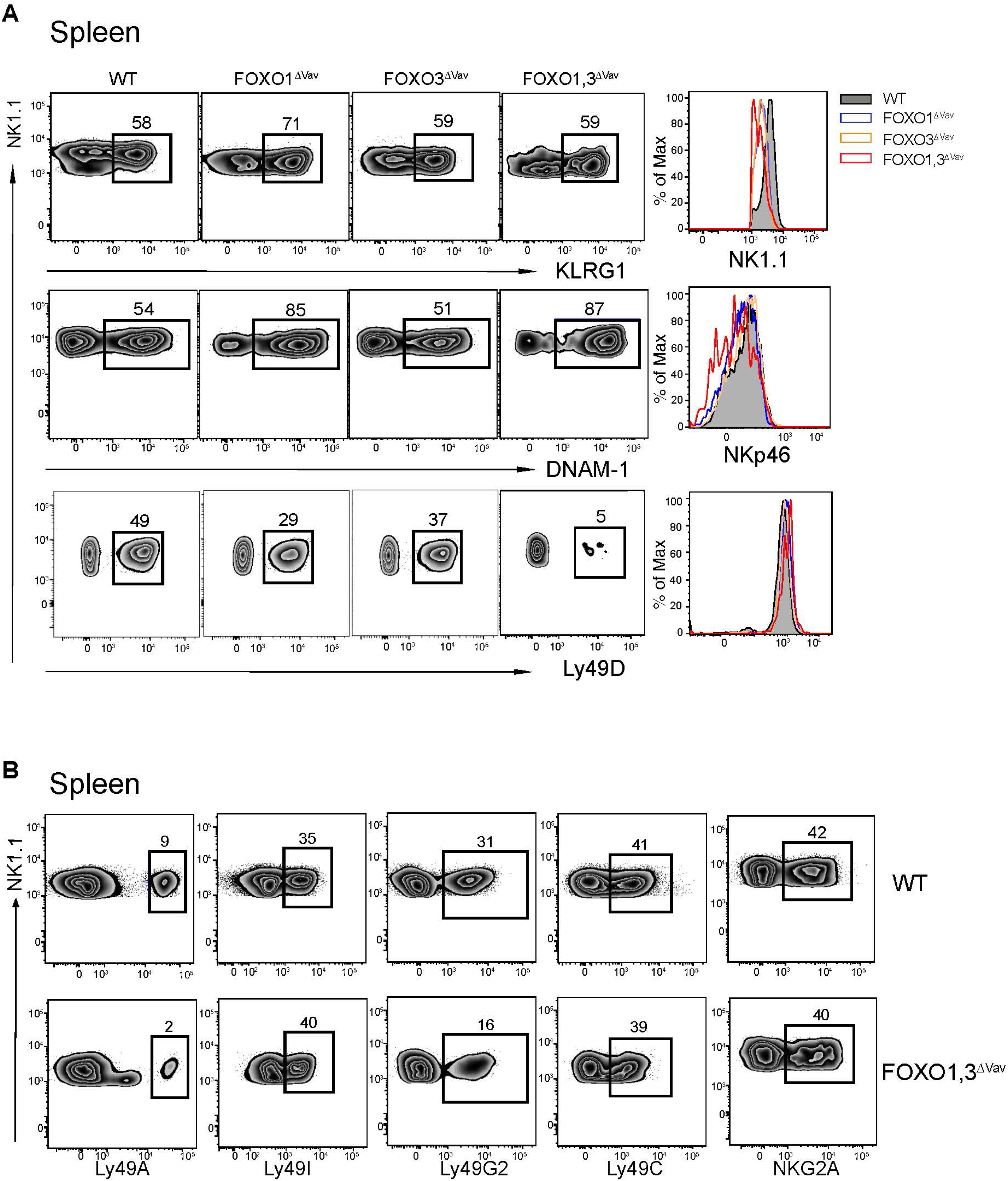
Splenic NK cells from FOXO1,3Δ^Vav^ have perturbed activating and inhibitory receptor expression. Representative flow cytometry plots showing activating (A) and inhibitory (B) receptor expression on splenic NK cells.

**Fig. S4.**
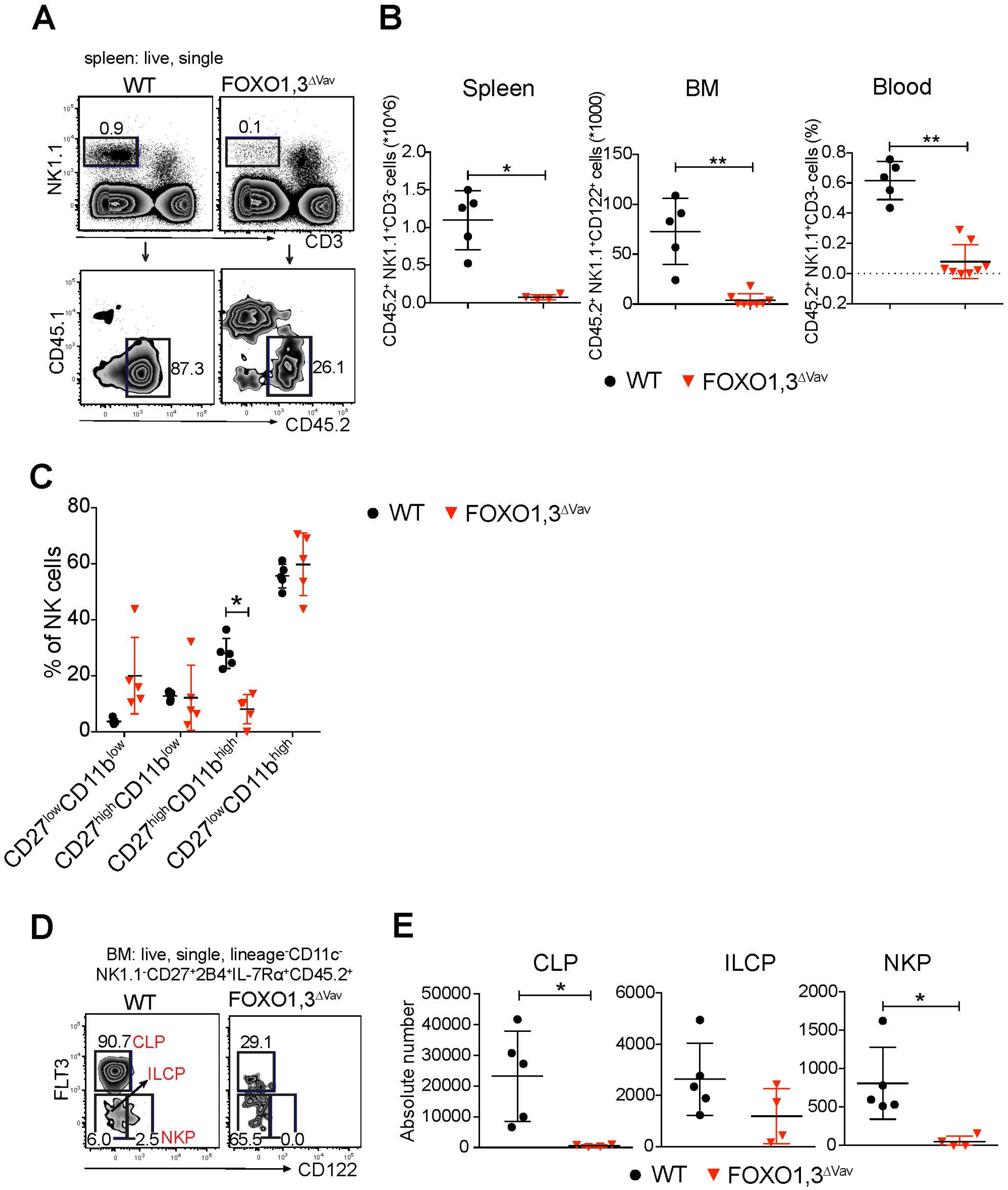
The NK defect in FOXO1,3Δ^Vav^ mice is intrinsic to hematopoiesis. (A) Representative flow cytometry profile showing NK cell reconstitution in spleen 12 weeks post-transplantation. CD45.1 recipient mice were irradiated and infused with unfractionated BM cells from CD45.2 donors with the indicated genotype. Reconstituted NK cells were defined as NK1.1^+^CD3^+^CD45.2^+^CD45.1^−^ cells. (B) Total number (in spleen and BM) and percentage (in blood) of donor NK cells 12 weeks post-transplantation (*n* = 4-8). (C) Frequency of splenic CD45.2+ NK cells at indicated maturation stages 12 weeks post-transplantation (*n* = 4-5). (D) Representative flow cytometry profiles showing the identification of CD45.2^+^ (donor) CLP, ILCP and NKP in the transplanted animals. (E) Total number of CLP, ILCP, and NKP in the transplanted animals (*n* = 4-5). In panels B, C and E: dots represent individual analyzed animals; p-values were calculated using Mann-Whitney; bars indicate mean and SD; *, ** and ** indicate p-values <0.05, <0.01 and <0.001 respectively.

**Fig. S5.**
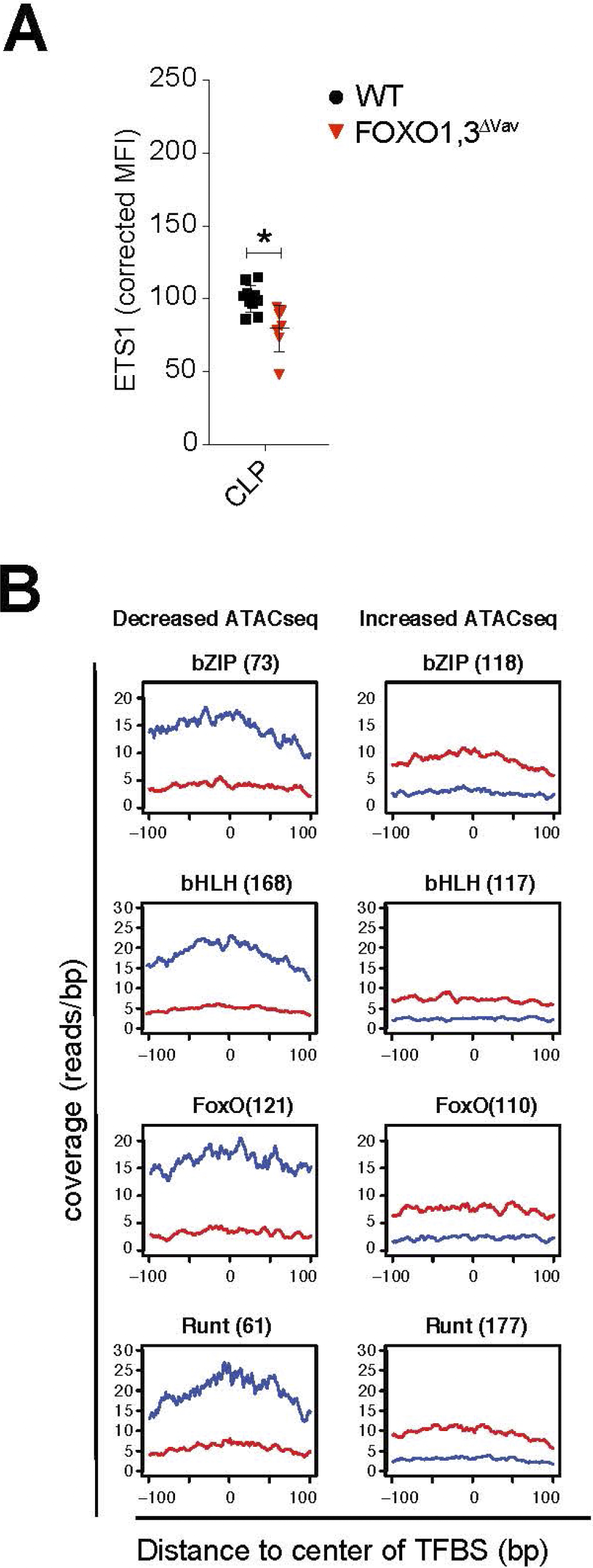
ETS1 protein expression and transcription factor cut-profiles. (A) ETS1 expression at protein level as measured by intracellular flow-cytometry. Dots represent individual analyzed animals; p-values were calculated using Mann-Whitney-Wilcoxon test; bars indicate mean and SD; * indicate p-values <0.05. (B) Cut-profile of differential ATAC-seq peaks with bZIP, bHLH, FoxO and Runt transcription factor binding sites (TFBS). The number of TFBS found within the differential peaks is indicated in brackets.

**Fig. S6.**
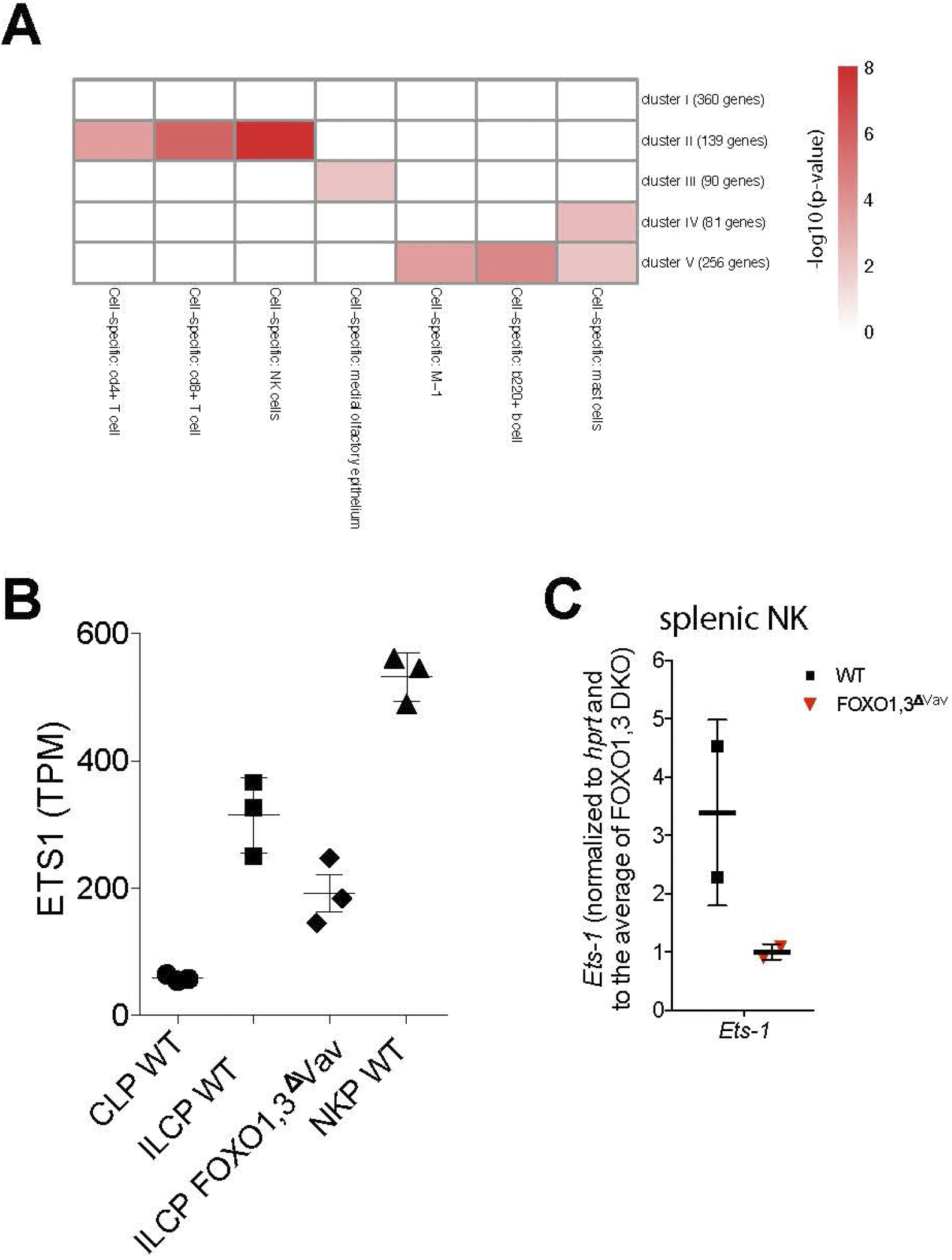
Annotation of developmentally regulated genes and comparison of FOXO expression. (A) Metascape gene enrichment analysis on the gene clusters from Figure 6D. (B) Ets1 expression (TPM) from RNAseq on CLP, ILCP and NKP from indicated genotype. (C) RTqPCR of Ets1 and Tbx21 from WT and FOXO1,3Δ^Vav^ NK cells from spleen.

